# Moving and static faces, bodies, objects and scenes are differentially represented across the three visual pathways

**DOI:** 10.1101/2022.11.30.518408

**Authors:** Emel Küçük, Matthew Foxwell, Daniel Kaiser, David Pitcher

## Abstract

Models of human cortex propose the existence of neuroanatomical pathways specialised for different behavioural functions. These pathways include a ventral pathway for object recognition, a dorsal pathway for performing visually guided physical actions and a recently proposed third pathway for social perception. In the current study we tested the hypothesis that different categories of moving stimuli are differentially processed across the dorsal and third pathways according to their behavioural implications. Human participants (*N*=30) were scanned with functional magnetic resonance imaging (fMRI) while viewing moving and static stimuli from five categories (faces, bodies, scenes, objects, and scrambled objects). Whole brain group analyses showed that moving bodies and moving objects increased neural responses in bilateral V5/MT+ and intraparietal sulcus (IPS), parts of the dorsal pathway. In addition, moving faces and moving bodies increased neural responses in bilateral V5/MT+ and the right posterior superior temporal sulcus (rpSTS), parts of the third pathway. This pattern of results was also supported by a separate region of interest (ROI) analysis showing that moving stimuli produced more robust neural responses for all visual object categories, particularly in lateral and dorsal brain areas. Our results suggest that dynamic naturalistic stimuli from different categories are routed along specific visual pathways that process their unique behavioural implications.

## Introduction

Explaining the neural processes that enable humans to perceive, understand and interact with the people, places, and objects we encounter in the world is a fundamental aim of visual neuroscience. An experimentally rich theoretical approach in pursuit of this goal has been to show that dissociable cognitive functions are performed in anatomically segregated cortical pathways. For example, influential models of the visual cortex propose it contains two functionally distinct pathways:. A ventral pathway specialised for visual object recognition, and a dorsal pathway specialised for performing visually guided physical actions (Kravitz et al., 2011; Kravitz et al., 2013; Milner & Goodale, 1995; Ungerleider & Mishkin, 1982). Despite the influence of these models, neither can account for the neural processes that underpin human social interaction. Social interactions are predicated on visually analysing and understanding the actions of others and responding appropriately. One region of the brain in particular, the superior temporal sulcus (STS), computes the sensory information that facilitates these processes (Allison et al., 2000; Kilner, 2011; Perrett et al., 1992). We recently proposed the existence of a visual pathway specialised for social perception (Pitcher & Ungerleider, 2021). This pathway projects from the primary visual cortex into the STS, via the motion-selective area V5/MT (Watson et al., 1993). The aim of the current study was to test a prediction of our model by contrasting the response to moving and static visual stimuli across the proposed three visual pathways. Specifically, we predicted moving biological stimuli (e.g., faces and bodies) are preferentially processed along a dedicated neural pathway that includes V5/MT and the STS, compared to moving stimuli of non-biological categories.

The STS selectively responds to moving biological stimuli (e.g., faces and bodies) and computes the visual social cues that help us understand and interpret the actions of other people. These include facial expressions (Phillips et al., 1997; Sliwinska, Elson, et al., 2020), eye gaze (Campbell et al., 1990; Pourtois et al., 2004; Puce et al., 1998), body movements (Beauchamp et al., 2003; Grossman & Blake, 2002) and the audio-visual integration of speech (Beauchamp et al., 2010; Calvert et al., 1997). However, the connectivity between early visual cortex and the STS remains poorly characterised. This led some researchers to view the STS as an extension of the ventral pathway, rather than as a functionally and anatomically independent pathway in its own right. For example, models of face processing propose that all facial aspects (e.g., identity and expression recognition) are processed using the same early visual mechanisms (Bruce & Young, 1986; Calder & Young, 2005; Haxby et al., 2000; Pitcher, Walsh, et al., 2011) before diverging at higher levels of processing, rather than as dissociable processes that begin in early visual cortex. Contrary to this view, alternate models propose that dynamic facial information is preferentially processed in a dissociable cortical pathway that projects from early visual cortex, via the motion-selective area V5/MT directly into the STS (Duchaine & Yovel, 2015; LaBar et al., 2003; O’Toole et al., 2002; Pitcher et al., 2014). This is consistent with our model of the third visual pathway which predicts that moving faces and bodies will selectively evoke neural activity in a pathway projecting from V1 to the STS, via V5/MT as shown in Figure 1 (Pitcher & Ungerleider, 2021).

**Figure 1.**
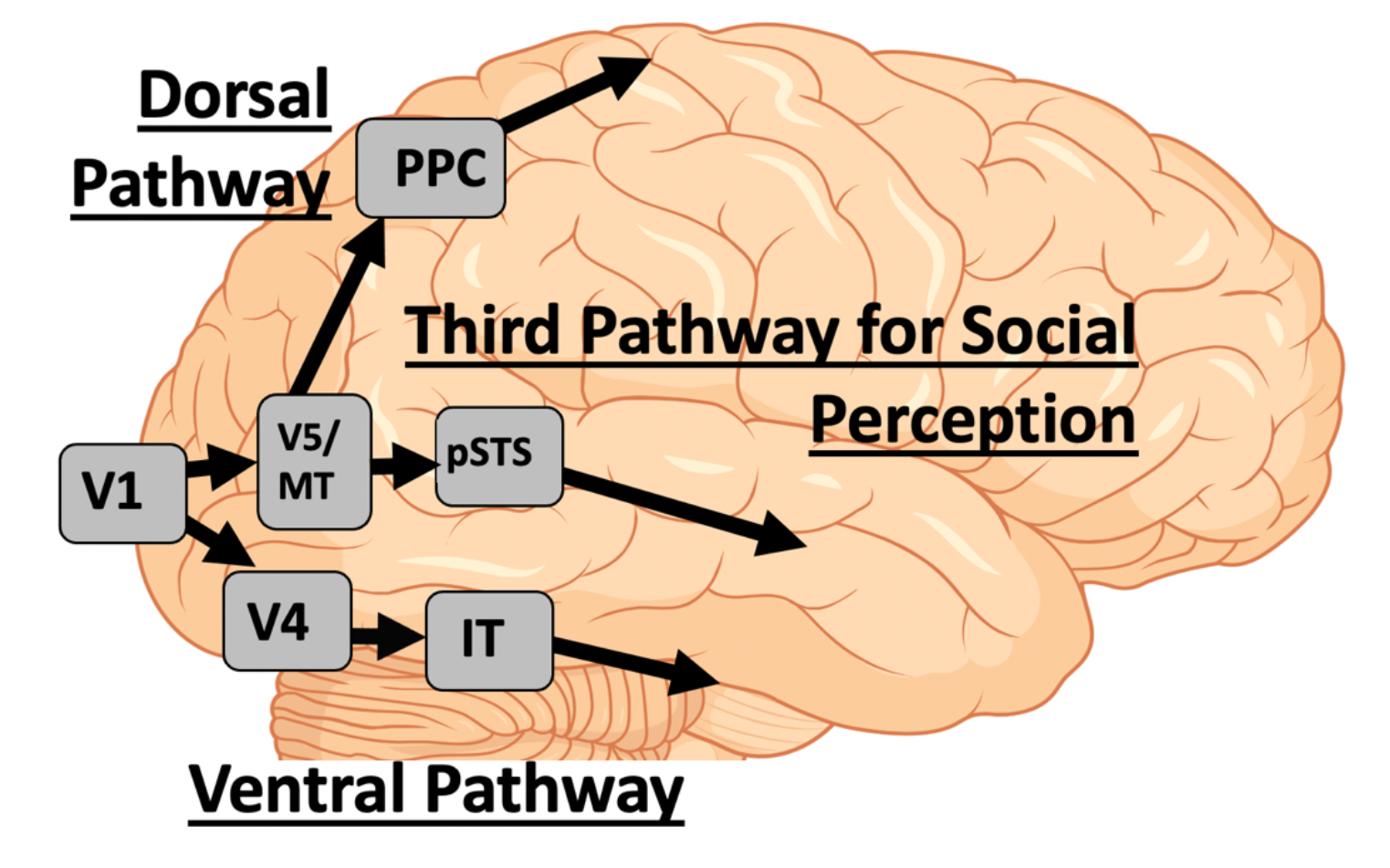
The three visual pathways (Pitcher & Ungerleider, 2021). The ventral pathway projects from V1 via ventral V4 into the inferior temporal (IT) and anterior inferior temporal cortex. The dorsal pathway projects from V1 to V5/MT into the posterior parietal cortex (PPC) and then to the motor cortex. The third visual pathway for social perception projects from V1 to V5/MT and then to the posterior STS (pSTS).

The cognitive and behavioural functions performed in a particular brain area can be deduced (at least partially) by the anatomical connectivity of that area (Boussaoud et al., 1990; Desimone & Ungerleider, 1986; Kravitz et al., 2011; Kravitz et al., 2013; Ungerleider & Desimone, 1986). This approach leads to hierarchical models that dissociate cognitive functions based on behavioural goals (e.g., visually recognising a friend, or reaching to shake their hand, or interpreting their mood). An alternate conceptual approach for studying the cognitive functionality of the brain has been to take a modular approach (Fodor, 1983). Modularity favours the view that cortex contains discrete cortical patches that respond to specific visual characteristics such as motion (Watson et al., 1993), or to stimulus categories including objects (Malach et al., 1995), faces (Kanwisher et al., 1997), scenes (Epstein & Kanwisher, 1998) and bodies (Downing et al., 2001). While there has sometimes been an inherent tension between anatomical and modular models of cortical organisation (Hein & Knight, 2008; Kanwisher, 2010), it has also been argued that different conceptual and methodical approaches can reveal cortical functionality at different levels of understanding (de Haan & Cowey, 2011; Grill-Spector & Malach, 2004; Walsh & Butler, 1996).

The aim of the present study was to measure the neural responses at both the hierarchical and modular level by manipulating the visual stimuli on two dimensions; moving versus static stimuli, or the object category of the stimuli (faces, bodies, scenes, and objects). Participants were scanned using functional magnetic resonance imaging (fMRI) while viewing 3-second videos or static images taken from the videos. The visual categories included were faces, bodies, objects, scenes, and scrambled objects (Figure 2). This set of stimuli can be used to identify a series of category-selective areas across the brain that include face areas (Haxby et al., 2000), body areas (Peelen & Downing, 2007), scene areas (Epstein, 2008) and object areas (Malach et al., 1995). We have previously used this design to functionally dissociate the neural response across face areas (Pitcher, Dilks, et al., 2011; Pitcher et al., 2014) and across the lateral and ventral surfaces of the occipitotemporal cortex (Pitcher et al., 2019). However, these prior studies lacked the necessary experimental conditions and the whole brain coverage to systematically compare the response to motion and visual category across the entire brain. The present study systematically compares the responses to different moving and static stimulus categories across the whole brain, as well as in targeted region-of-interest analyses. This provides detailed insights into how dynamically presented visual categories are routed along the three visual pathways.

**Figure 2.** Examples of the static images taken from the 3-second movie clips depicting faces, bodies, scenes, objects, and scrambled objects. Still images taken from the beginning, middle and end of the corresponding movie clip.

## Materials and Methods

### Participants

A total of thirty participants (20 females; age range 18 to 48 years old; mean age 23 years) with normal, or corrected-to-normal, vision gave informed consent as directed by the Ethics committee at the University of York. Data from 24 participants was collected for a previous fMRI experiment (Nikel et al., 2022) and re-analysed for the current study.

### Stimuli

Dynamic stimuli were 3-second movie clips of faces, bodies, scenes, objects and scrambled objects designed to localise the category-selective brain areas of interest (Pitcher, Dilks, et al., 2011). There were sixty movie clips for each category in which distinct exemplars appeared multiple times. Movies of faces and bodies were filmed on a black background, and framed close-up to reveal only the faces or bodies of 7 children as they danced or played with toys or adults (who were out of frame). Fifteen different locations were used for the scene stimuli which were mostly pastoral scenes shot from a car window while driving slowly through leafy suburbs, along with some other films taken while flying through canyons or walking through tunnels that were included for variety. Fifteen different moving objects were selected that minimised any suggestion of animacy of the object itself or of a hidden actor pushing the object (these included mobiles, windup toys, toy planes and tractors, balls rolling down sloped inclines). Scrambled objects were constructed by dividing each object movie clip into a 15 by 15 box grid and spatially rearranging the location of each of the resulting movie frames. Within each block, stimuli were randomly selected from within the entire set for that stimulus category (faces, bodies, scenes, objects, scrambled objects). This meant that the same actor, scene or object could appear within the same block but given the number of stimuli this did not occur regularly.

Static stimuli were identical in design to the dynamic stimuli except that in place of each 3-second movie we presented three different still images taken from the beginning, middle and end of the corresponding movie clip. Each image was presented for one second with no ISI, to equate the total presentation time with the corresponding dynamic movie clip (Figure 2). This same stimulus set has been used in our prior fMRI studies of category-selective areas (Handwerker et al., 2020; Sliwinska et al., 2022).

#### Procedure and Data Acquisition

Functional data were acquired over 8 blocked-design functional runs lasting 234 seconds each. Each functional run contained three 18-second rest blocks, at the beginning, middle, and end of the run, during which a series of six uniform colour fields were presented for three seconds. Participants were instructed to watch the movies and static images but were not asked to perform any overt task.

Functional runs presented either movie clips (four dynamic runs) or sets of static images taken from the same movies (four static runs). For the dynamic runs, each 18-second block contained six 3-second movie clips from that category. For the static runs, each 18-second block contained 18 one-second still snapshots, composed of six triplets of snapshots taken at one second intervals from the same movie clip. Dynamic / static runs were run in the following order: 2 dynamic, 2 static, 2 dynamic, 2 static.

Imaging data were acquired using a 3T Siemens Magnetom Prisma MRI scanner (Siemens Healthcare, Erlangen, Germany) at the University of York. Functional images were acquired with a twenty-channel phased array head coil and a gradient-echo EPI sequence (38 interleaved slices, repetition time (TR) = 3 sec, echo time (TE) = 30ms, flip angle = 90°; voxel size 3mm isotropic; matrix size = 128 × 128) providing whole brain coverage. Slices were aligned with the anterior to posterior commissure line. Structural images were acquired using the same head coil and a high-resolution T-1 weighted 3D fast spoiled gradient (SPGR) sequence (176 interleaved slices, repetition time (TR) = 7.8 sec, echo time (TE) = 3ms, flip angle = 20°; voxel size 1mm isotropic; matrix size = 256 × 256).

#### Data and Materials Availability

Data for all participants is available on requests. The whole brain group activation maps used in generate the images in Figures 3, 4 and 5 are available at https://osf.io/3w4ps/

**Figure 3.**
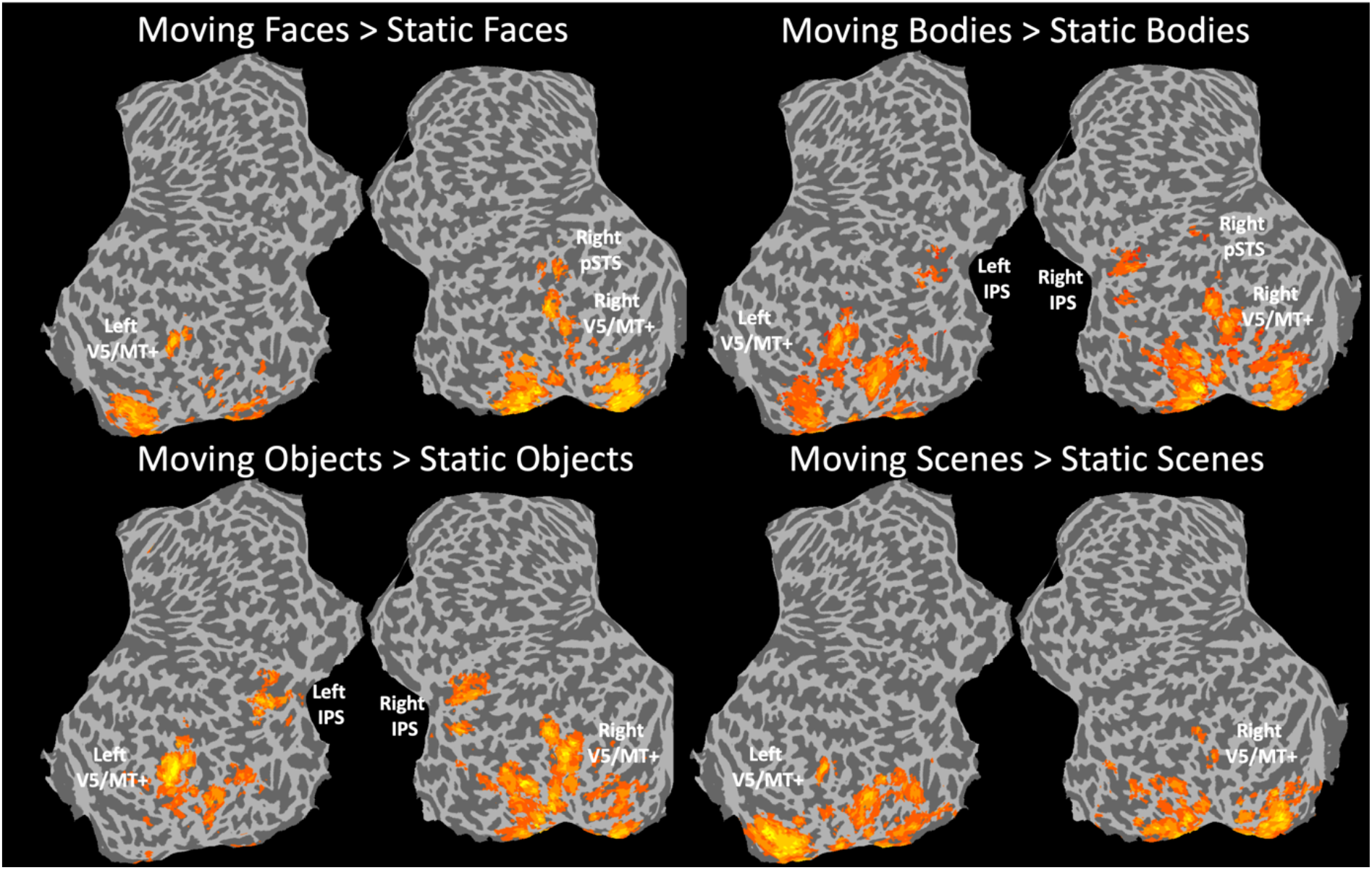
Results of the group whole brain contrasts showing moving greater than static neural activity for each visual category (faces, bodies, objects, and scenes). The moving > static faces and moving > static bodies contrasts produced clusters in the posterior STS, part of the third visual pathway. The moving > static bodies and moving > static objects contrasts produced clusters in the bilateral IPS, part of the dorsal visual pathway. None of the contrasts yielded significant clusters in the ventral pathway. Maps generated using a t-statistical threshold of *p* = 0.001 and a cluster correction of 50 contiguous voxels.

**Figure 4.**
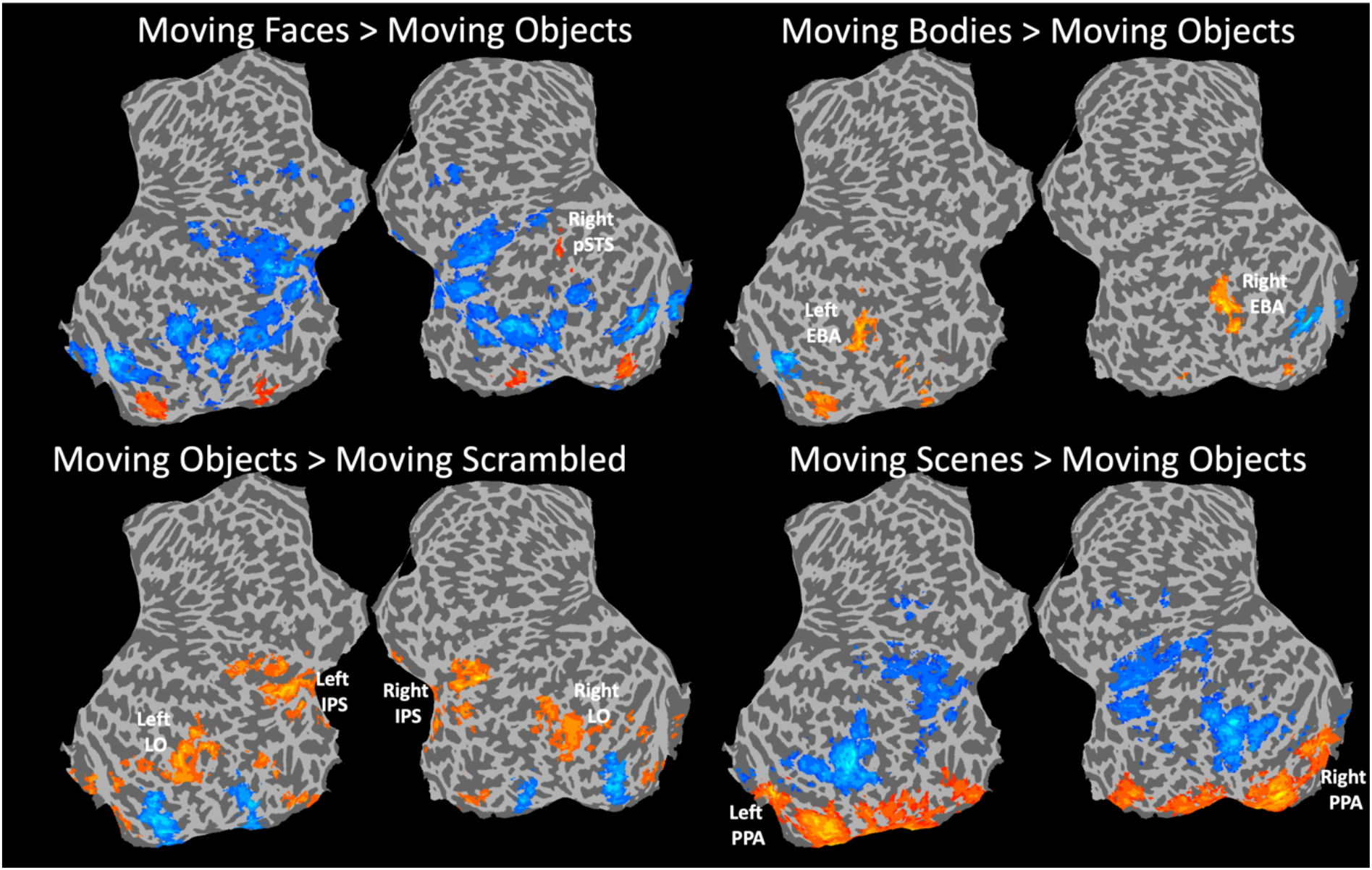
Results of the group whole brain contrasts showing category-selective areas identified using moving stimuli. Moving faces > moving objects produced a significant cluster in the right pSTS. Moving bodies > moving objects produced significant clusters in the bilateral EBA. Moving objects > moving scrambled objects produced a significant cluster in the bilateral LO and IPS. Moving scenes > moving objects produced significant clusters in the bilateral PPA. None of the contrasts yielded significant clusters in the ventral pathway. Maps generated using a t-statistical threshold of *p* = 0.001 and a cluster correction of 50 contiguous voxels.

**Figure 5.**
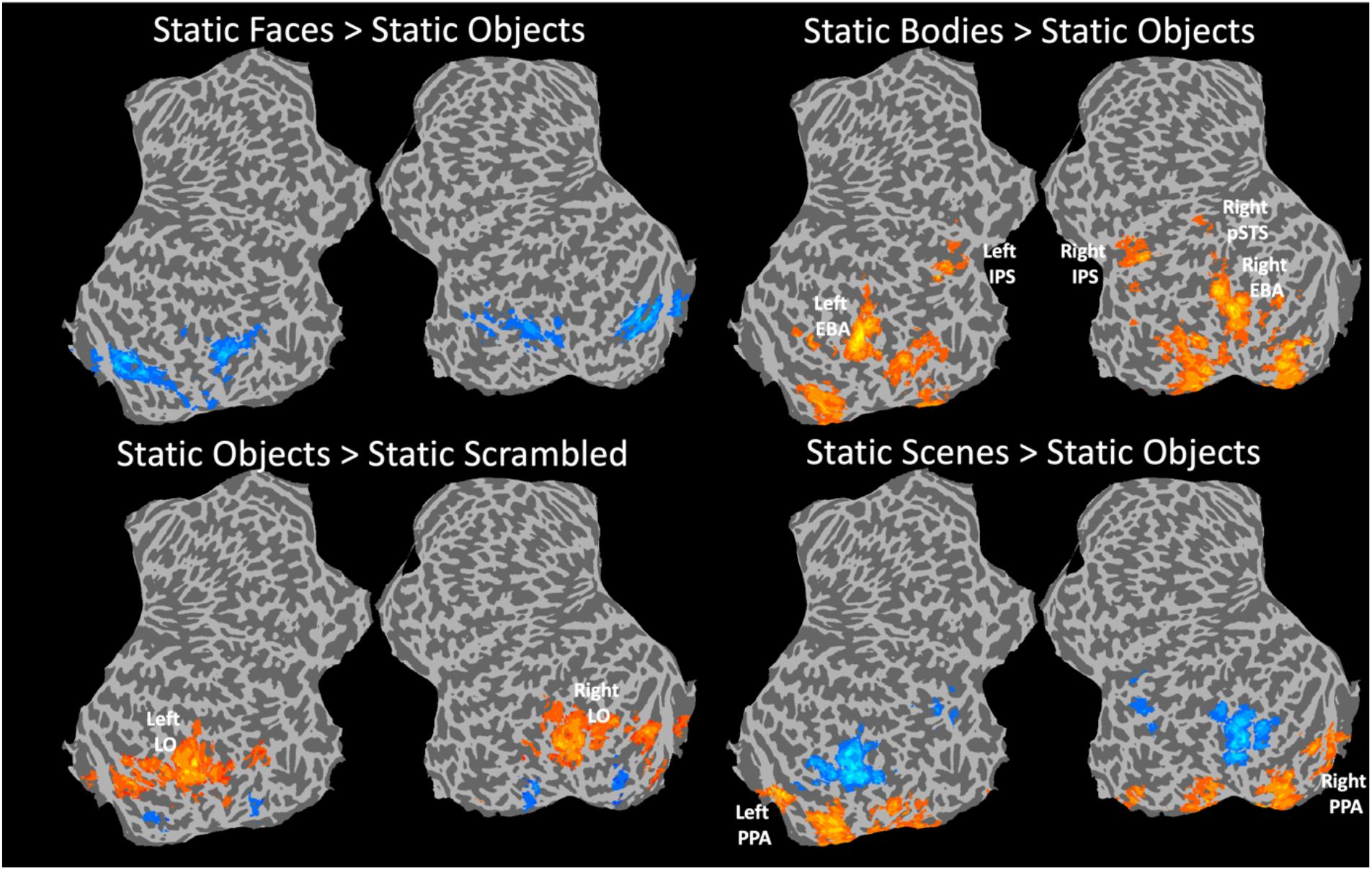
Results of the group whole brain contrasts showing category-selective areas identified using static stimuli. Static faces > static objects produced no significant clusters. Static bodies > static objects produced significant clusters in the bilateral EBA, the right pSTS and bilateral IPS. Static objects > static scrambled objects produced a significant cluster in the bilateral LO and IPS. Static scenes > static objects produced significant clusters in the bilateral PPA. None of the contrasts yielded significant clusters in the ventral pathway. Maps generated using a t-statistical threshold of *p* = 0.001 and a cluster correction of 50 contiguous voxels.

#### Imaging Analysis

Functional MRI data were analysed using AFNI (http://afni.nimh.nih.gov/afni). Images were slice-time corrected and realigned to the third volume of the first functional run and to the corresponding anatomical scan. All data were motion corrected and any TRs in which a participant moved more than 0.3mm in relation to the previous TR were discarded from further analysis. The volume-registered data were spatially smoothed with a 4-mm full-width-half-maximum Gaussian kernel. Signal intensity was normalised to the mean signal value within each run and multiplied by 100 so that the data represented percent signal change from the mean signal value before analysis.

Data from all runs were entered into a general linear model (GLM) by convolving the standard hemodynamic response function with the regressors of interest (faces, bodies, scenes, objects, and scrambled objects) for dynamic and static functional runs. Regressors of no interest (e.g., 6 head movement parameters obtained during volume registration and AFNI’s baseline estimates) were also included in the GLM. Data from all thirty participants were entered into a group whole brain analysis. Group whole brain contrasts were generated to quantify the neural responses across three experimental conditions (see results section). Activation maps were calculated using a t-statistical threshold of *p* = 0.001 and a cluster correction of 50 contiguous voxels. For display purposes, volumetric whole brain maps were projected onto a flat cortical surface using SUMA (https://afni.nimh.nih.gov/Suma).

Data for all participants were also individually analysed to localise the regions of interest (ROIs) using data collected in the four dynamic runs. Category selective ROIs were identified by generating significance maps of the brain using an uncorrected statistical threshold of *p* = 0.001. Face-selective ROIs were identified for each participant using a contrast of greater activation evoked by faces than by objects. Body-selective ROIs were identified for each participant using a contrast of greater activation evoked by bodies than by objects. Object-selective ROIs were identified for each participant using a contrast of greater activation evoked by objects than by scrambled objects. Scene-selective ROIs were identified for each participant using a contrast of greater activation evoked by scenes than by objects.

## Results

### Group Whole Brain Analysis: Moving > Static within category

Data from all thirty participants were entered into a group whole brain ANOVA to generate the contrasts of interest. The first analysis established the neural differences between moving and static stimuli using the following contrasts: moving faces > static faces; moving bodies > static bodies; moving objects > static objects; moving scenes > static scenes (Figure 3). Results were consistent with the predicted differential responses across the three visual pathways. All contrasts generated clusters beginning in early visual cortex and extending into V5/MT+ as defined by probabilistic maps for combining functional imaging data with cytoarchitectonic maps (Eickhoff et al., 2005). This was to be expected as the moving stimuli generated more visual energy than static stimuli resulting in greater activation across early visual areas (Snowden et al., 1991).

Dissociations between different object categories were revealed in higher visual areas. Moving faces greater than static faces resulted in clusters in the right posterior STS (in the third pathway), but not in the fusiform gyrus (in the ventral pathway). Moving bodies greater than static bodies resulted in clusters in the right posterior STS (in the third pathway) and the bilateral intraparietal sulcus (IPS) (in the dorsal pathway). This right lateralized response to faces in the STS is consistent with our prior TMS evidence (Sliwinska & Pitcher, 2018). The large contiguous region encompassing V5/MT+ also extended into the expected location of the extrastriate body area (EBA) bilaterally. This is consistent with studies showing the two areas spatially overlap (Downing et al., 2007). Moving objects greater than static objects resulted in clusters in the bilateral IPS (in the dorsal pathway). Similar to the bodies contrast, the large contiguous region encompassing V5/MT+ also extended into the expected location of the lateral occipital cortex (LO) bilaterally. The moving scenes greater than static scenes contrast did not produce any significant clusters in the parahippocampal gyrus (part of the ventral pathway).

### Group Whole Brain Analysis: Moving stimuli between categories

The second analysis investigated the category-selective neural responses to the four different visual stimulus categories (faces, bodies, objects, and scenes) using moving stimuli. Group whole brain contrasts were calculated for faces (moving faces > moving objects), bodies (moving bodies > moving objects), objects (moving objects > moving scrambled objects) and scenes (moving scenes > moving objects) (Figure 4). Results were partially consistent with prior fMRI studies of high-level category-selective visual areas. Moving faces > moving objects produced a cluster in the right pSTS. Notably, this analysis did not reveal face-selective regions in the ventral stream, such as the fusiform face area (FFA), presumably due to inter-individual variation in the anatomical location of the region. These regions were, however, robustly identified in a participant-specific region-on-interest analysis (see below). Moving bodies > moving objects produced a cluster in the bilateral EBA. Moving objects > moving scrambled objects produced a cluster in the bilateral LO and bilateral IPS. Moving scenes > moving objects produced a cluster in the bilateral parahippocampal place area (PPA). All four contrasts also produced significant clusters in early visual cortex, a result consistent with prior studies showing greater visual motion produces greater neural activity in the primary visual cortex (Snowden et al., 1991).

### Group Whole Brain Analysis: Static stimuli between categories

The third analysis investigated the category-selective neural responses to the four different visual stimulus categories (faces, bodies, objects, and scenes) using static stimuli. Group whole brain contrasts were calculated for faces (static faces > static objects), bodies (static bodies > static objects), objects (static objects > static scrambled objects) and scenes (static scenes > static objects) (Figure 5). Results were partially consistent with prior fMRI studies of high-level category-selective visual areas. Static faces > static objects produced no significant clusters, but face-selective clusters were again reliably localised in a participant-specific region-of-interest analysis (see below). Static bodies > static objects produced clusters in the bilateral EBA, right pSTS and bilateral IPS. Static objects > static scrambled objects produced clusters in the bilateral LO. Static scenes > static objects produced clusters in the bilateral PPA. A detailed comparison of the regions identified with moving and static stimuli is provided in the following region-of-interest analysis.

### Region of Interest (ROI) Analysis

Finally, we evaluated the use of moving stimuli for functionally localising category-selective regions in the three visual pathways. We analysed data for all participants individually to separately localise the regions of interest (ROIs) using dynamic and static stimuli for all four stimulus categories (faces, scenes, bodies, and objects). Significance maps were calculated for each participant individually, using an uncorrected statistical threshold of *p* = 0.001 for all four contrasts of interest. If the ROI was present in that participant, we identified the MNI co-ordinates of the peak voxel and the number of contiguous voxels in the ROI. Mean results are displayed in Table 1.

**Table 1.**
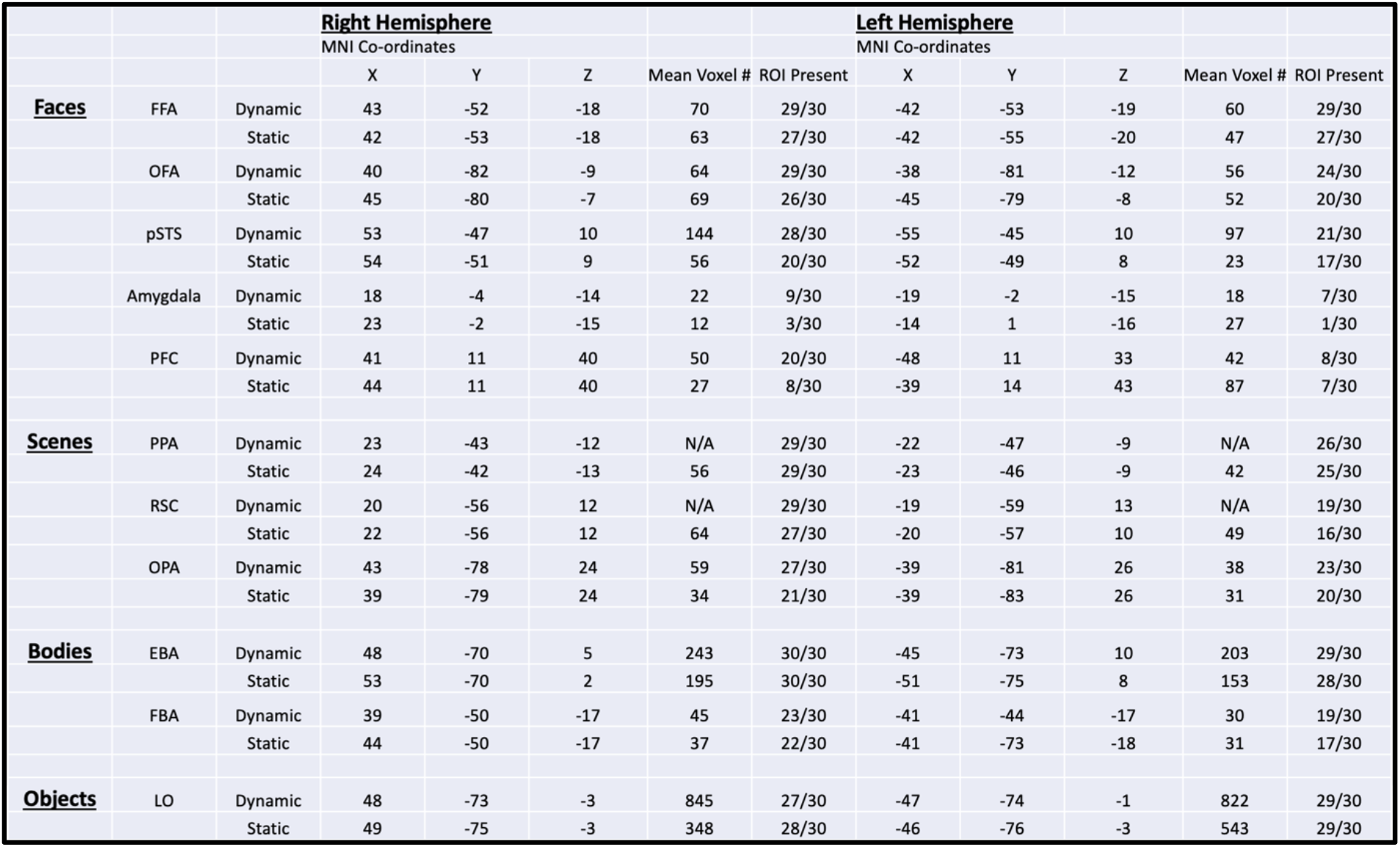
The results of the ROI analysis for dynamic and static stimuli for face, scene, body, and object areas in 30 participants. Face-selective areas were more robustly identified with dynamic stimuli in the pSTS and prefrontal cortex (PFC). A contrast of dynamic scenes > dynamic objects produced large clusters encompassing the PPA, retrosplenial cortex (RSC) and large areas of the visual cortex in 26 participants. The object-selective LO also produced large clusters contiguous with early visual areas when defined using a contrast of moving objects > scrambled objects. The body-selective EBA was the most consistently identified ROI across participants, regardless of whether it was identified using dynamic or static stimuli.

Face-selective ROIs were identified in two separate analyses, using a contrast of moving faces greater than moving objects and a contrast of static faces greater than static objects. We attempted to localise face-selective voxels in five commonly studied face ROIs. These were the fusiform face area (FFA), occipital face area (OFA), pSTS, the amygdala and in the prefrontal cortex (PFC). Results showed that face ROIs were present in more participants in the right hemisphere. The pSTS and PFC were identified in more participants when defined using moving than static stimuli. The pSTS and PFC ROIs were also larger when identified with moving stimuli. These results are consistent with some prior studies (Fox et al., 2009; Nikel et al., 2022; Pitcher, Dilks, et al., 2011; Pitcher et al., 2019) but it is important to note that other studies have demonstrated that the FFA can also exhibit a greater response to moving faces than static faces (Pilz et al., 2011; Schultz & Pilz, 2009).

Scene-selective ROIs were identified in two separate analyses, using a contrast of moving scenes greater than moving objects and a contrast of static scenes greater than static objects. We attempted to localise scene-selective voxels in the three commonly studied scene-selective ROIs. These were the parahippocampal place area (PPA), retrosplenial cortex (RSC) and the occipital place area (OPA). Results demonstrated that moving scenes greater than moving objects generated activations in the PPA and RSC that were contiguous not only with each other but also with large sections of visual cortex in twenty-three participants. These large clusters yielded a large number of contiguous voxels, so that efficient functional localisation needs to also rely on anatomical constraints or spatial constraints from existing group templates. The OPA was spatially distinct from this cluster in most participants (see Table 1). Static scenes greater than static objects demonstrated results more consistent with earlier fMRI studies of scene-selective ROIs. The PPA, RSC and OPA were successfully localised in the right hemisphere of most participants. These ROIs were less consistent in the left hemisphere, notably the RSC (see Table 1).

Body-selective ROIs were identified in two separate analyses, using a contrast of moving bodies greater than moving objects and a contrast of static bodies greater than static objects. We attempted to localise the two most studied body-selective ROIs, the extrastriate body area (EBA) and fusiform body area (FBA). Notably, the EBA was identified bilaterally across more participants using both moving and static contrasts than any other category-selective ROI (Table 1). Results further showed that the EBA was larger when defined using moving than static stimuli, but this was not the case with the FBA. This is consistent with prior evidence showing that lateral category-selective brain areas exhibit a greater response to moving stimuli more than static stimuli (Pitcher et al., 2019).

Finally, we defined the object-selective area LO using contrasts of moving objects > moving scrambled objects and a contrast of static objects > static scrambled objects. Results showed a similar pattern to the scene-selective ROIs, namely that defining LO using moving stimuli produced large bilateral ROIs that were contiguous with early visual cortex (Table 1). By contrast, defining LO using static stimuli was more consistent with prior studies (Malach et al., 1995; Pitcher et al., 2009).

## Discussion

In the present study we used fMRI to measure the neural responses to dynamic and static stimuli from different visual categories (faces, bodies, scenes, and objects). Our aim was to establish the brain areas that selectively respond to moving stimuli of different categories across the three visual pathways. Results supported functional dissociations consistent with the recently proposed third visual pathway for social perception (Pitcher & Ungerleider, 2021). Specifically, contrasts of moving faces greater than static faces and moving bodies greater than static bodies produced clusters in bilateral V5/MT+ and in the right posterior STS (Figure 3). Moving objects greater than static objects also produced clusters in bilateral intraparietal sulcus (IPS), part of the dorsal visual pathway for visually guided action (Milner & Goodale, 1995). Interestingly, moving bodies greater than static bodies activated both pathways, with greater activation for the moving stimuli in both STS and IPS. These results demonstrate that the motion of stimuli from high-level object categories is preferentially processed in the lateral and dorsal areas of the visual cortex, more than category-selective areas on the ventral brain surface. These results also suggest that there is a critical division between biological- and non-biological stimuli across the dorsal and third visual pathways.

Preferential representations of dynamic face stimuli in the STS aligns with the role of the STS in social perception (Allison et al., 2000; Kilner, 2011; Perrett et al., 1992) and representing aspects of the face that can change rapidly such as expression, gaze and mouth movements (Haxby et al., 2000). Accordingly, selectivity to facial motion facilitates emotion perception in facial expressions and bodies (Atkinson et al., 2004; Kilts et al., 2003) and audio-visual integration of speech (Young et al., 2020). The STS has also been implicated in biological motion perception, producing a greater response to motion stimuli depicting jumping, kicking, running and throwing movements than control motion (Grossman et al., 2000). Such motion stimuli, as well as changeable aspects of the face, convey information that may provoke attributions of intentionality and personality of other individuals (Adolphs, 2002). Moreover, previous studies have shown that the STS exhibits high selectivity to social in contrast to non-social stimuli (Lahnakoski et al., 2012; Watson et al., 2014). In addition to finding that the STS seemed to be ‘people selective’, Watson et al. (2014) demonstrated the multisensory nature of the STS, proposing this region plays a vital role in combining socially relevant information across modalities. This discrimination between social and non-social stimuli is consistent with our proposal of a third pathway connecting V5 and STS, which is specialised for processing dynamic aspects of social perception (Pitcher & Ungerleider, 2021).

Our results for moving bodies are consistent with previous studies of body-selectivity in humans, which show that the EBA and the FBA respond more strongly to human bodies and body parts than faces, objects, scenes and other stimuli (Downing et al., 2001; Peelen & Downing, 2005, 2007). The posterior superior temporal sulcus (pSTS) has also been implicated in the perception of biological motion through faces or bodies (Grossman et al., 2000; Puce & Perrett, 2003; Saygin, 2007). Previous studies have highlighted the dissociation in the response to dynamic and static presentation of bodies in lateral and ventral regions (Grosbras et al., 2012; Pitcher et al., 2019) that we also report here. The location of these body-selective regions in distinct neuroanatomical pathways, with the FBA located on the ventral surface (Peelen & Downing, 2005; Schwarzlose et al., 2005) and the EBA (Downing et al., 2001) and pSTS (Grossman et al., 2000) on the lateral surface, is consistent with these regions having different response profiles to static and dynamic images of bodies. Interestingly, we also observed significant clusters to body stimuli in the IPS, a brain area that is part of the dorsal processing stream for computing visually guided physical actions (Milner & Goodale, 1995). These clusters were present in the moving bodies greater than static bodies and static bodies greater than static objects contrasts, demonstrating that actual body motion and static bodies with implied motion are processed in the IPS (this was also true for the EBA). However, the moving bodies greater than moving objects contrast did not result in significant clusters in the IPS. This was because moving objects (but not static objects) are also processed in the IPS, so the responses to moving bodies and moving objects cancelled each other out. This pattern of results suggests that the multi-faceted behavioural relevance of bodies triggers processing in both the dorsal and third visual pathways: Bodies are not only relevant for inferring social information about others (just like faces), they also are critical for perceiving and evaluating visually guiding action (just like objects). Our results are therefore consistent with the idea that the differential routing of categorical information across the dorsal and third pathways is not determined by movement per se, but by the behavioural implications carried by the movement for a specific stimulus category.

Neuroimaging studies have identified multiple scene-selective brain regions in humans, including the parahippocampal place area (PPA) (Epstein & Kanwisher, 1998), the retrosplenial cortex (RSC) (Maguire et al., 2001) and the occipital place area (OPA) (Dilks et al., 2013). Previous studies have highlighted the OPA showing a greater response to dynamic than static scenes, whereas the PPA and RSC showed similar responses to dynamic and static scenes (Kamps et al., 2016; Korkmaz Hacialihafiz & Bartels, 2015; Pitcher et al., 2019). This is consistent with our result that moving scenes do not activate scene-selective areas in the ventral stream more strongly than static scenes. Additionally, this selective response in the OPA to dynamic scenes is seemingly not as a result of low-level information processing or domain-general motion sensitivity (Kamps et al., 2016). The way these scene-selective regions can be dissociated based on motion sensitivity aligns with their possible roles in scene processing. Mirroring the role of the ventral pathway in recognition and the dorsal pathway in visually guided action (Milner & Goodale, 1995), these findings are consistent with the hypothesis of two distinct scene processing systems engaged in navigation and other aspects of scene processing such as scene categorisation (Dilks et al., 2011; Persichetti & Dilks, 2016). The anatomical position of the OPA within the dorsal pathway is compatible with its motion sensitivity and role in visually guided navigation (Kamps et al., 2016). Whereas the ventral/medial location of the PPA and RSC aligns with their demonstration of less sensitivity to motion and role in other aspects of navigation and scene recognition. Notably, our study revealed much more widespread scene-selective clusters when localisation was performed with dynamic stimuli. These larger clusters may reflect the type of movement present in the scene stimuli: Rather than local movement of a foreground object (as present in the faces, body, and object stimuli), scene stimuli were characterised by more global movement patterns, where instead of the scene itself, the camera would move. Such global movement patterns may indeed be a relevant source of information in scene processing, where temporal dynamics in the information are often a consequence of the observer moving through the world. However, future studies need to systematically compare such global movements with scenes in which many local elements actively move (like trees and leaves move during a windy day) to delineate whether there are different consequences of these movement types on scene representation.

In addition to the group whole brain analyses we also performed ROI analyses for all four stimulus categories at the individual participant level (Table 1), allowing us to quantify whether moving stimuli can improve the quality of functional localization in the visual system. Prior fMRI studies have mostly used static images of stimuli from these categories to identify the relevant category-selective brain areas. However, a subset of areas are known to exhibit a greater neural response to moving more than static images from the preferred visual object category. These include face-selective areas in the superior temporal sulcus (Fox et al., 2009; LaBar et al., 2003; Pitcher, Dilks, et al., 2011; Puce et al., 1998), the scene-selective OPA in the transverse occipital sulcus (Kamps et al., 2016) and the body-selective EBA in the lateral occipital lobe (Pitcher et al., 2019). This spatial dissociation between moving and static stimuli was also observed in the region of interest (ROI) analyses performed for each individual participant (Table 1). Results were consistent with prior studies of these same areas (Pitcher, Dilks, et al., 2011; Pitcher et al., 2019; Sliwinska, Bearpark, et al., 2020). Our ROI results also demonstrate that moving stimuli successfully identify more ROIs across participants and larger ROIs than static stimuli for face, scene and body areas, particularly in lateral brain areas. By contrast, moving stimuli did not lead to systematic shifts in ROI peak coordinates, showing that the activations indeed stem from the same cortical areas. The differential selectivity for motion is believed to relate to the different cognitive functions performed on stimuli within the relevant category. For example, facial identity or facial expression recognition (Haxby et al., 2000) or scene recognition or spatial navigation (Kamps et al., 2016). From a methodological perspective, the results shown in Table 1 also demonstrate that the use of moving stimuli for fMRI functional localisers will more robustly identify category-selective ROIs for all four stimulus categories across both hemispheres. The choice of stimuli in functional localiser experiments depends on multiple factors, including how these regions are probed in the subsequent experiments. Our study, however, provides a valuable benchmark for ROI-based fMRI studies in the future, where functional localisation will be robustly achieved in a greater percentage of participants when moving stimuli are used instead of static stimuli.

## Acknowledgements

Thanks to Nancy Kanwisher for providing experimental stimuli.

